# The *Amt2* Gene is Key for *Cryptococcus neoformans* Transmigration Across The Blood-Brain Barrier and Closely Linked to Its Capsule Formation

**DOI:** 10.64898/2026.01.30.702728

**Authors:** Sainamthip Rangdist, Ekkasit Kanklang, Natchaya Pakdeesiriwong, Watsana Penkhrue, Chompunoot Wangboon, Oratai Weeranantanapan, Sirilak Chumkiew, Pathanin Chantree, Pongsakorn Martviset, Methee Chayakulkeeree, Mantana Jamklang

## Abstract

The cryptococcal *Amt* family of ammonium transporters have been identified from our previous studies as one of the most highly upregulated proteins during transmigration in an in vitro blood-brain barrier (BBB) model, however, the role of this gene family has never been reported. Therefore, this study aimed to investigate the role of the *Amt2* gene in the transmigration of *C. neoformans* across the BBB, examine its association with other common virulence factors, and assess its relevance to morphological changes in *C. neoformans.* The results showed that the *C. neoformans* mutant strain lacking the *Amt2* gene (*amt2*Δ) exhibited a significantly reduced ability to transmigrate across the BBB in an in vitro model. Our findings suggest that *C. neoformans* primarily utilizes a transcellular mechanism for invasion, as indicated by the FITC-dextran permeability assays. Additionally, the size of polysaccharide capsule was significantly smaller in the mutant strain compared to the wild-type. In conclusion, our study proposed that the *Amt2* gene plays a crucial role in both the transmigration process and capsule production in *C. neoformans*, without affecting morphological changes. Our study provides a foundation for future research into the underlying mechanisms of the *Amt2* gene in *C. neoformans* pathogenesis.

**Author summary:** *Cryptococcus neoformans* transmigrates the blood-brain barrier through various mechanisms, with transcellular migration being the major route leading to cryptococcal meningitis. In this study, we identified the *Amt2* gene, a member of the Amt family of ammonium transporters, as playing a crucial role in the fungus’s transmigration process. Our findings indicate that the *Amt2* gene promotes capsule production and facilitates the transmigration of *C. neoformans*, all while not causing damage to human endothelial cells.

## Introduction

*Cryptococcus* spp. is a pathogenic encapsulated yeast that causes life-threatening fungal invasion of the respiratory and central nervous systems (CNS), especially in immunocompromised patients (1, 2). In 2020, the global incidence of cryptococcosis was estimated at approximately 152,000 cases, leading to around 112,000 deaths yearly (3). This alert has become a significant concern for the international health systems. The two species most commonly associated with cryptococcosis in humans are *C. neoformans* and *C. gattii*. *C. neoformans* is globally distributed and colonized in a variety of biological niches including soil, pigeon droppings and decaying trees whereas *C. gattii* is specifically found in a tree species associated with tropical and subtropical climates (4, 5). Infections occur when inhaling dehydrated spores, leading to pulmonary cryptococcosis, which can subsequently disseminate to the CNS and cause cryptococcal meningitis (6, 7). *Cryptococcus* can infect the meninges and brain parenchyma, leading to brain lesions and cryptococcal meningoencephalitis (8). Focus on *C. neoformans*, the rationale behind the profound invasiveness of *C. neoformans* lies in its key virulence factors, which help it evade and resist host immunity. These factors include its polysaccharide capsule, melanin production, and thermotolerance. Additionally, *C. neoformans* produces urease and phospholipase B, which facilitate brain invasion and damage endothelial cells in the brain (9, 10). These factors contribute to *C. neoformans* being a major cause of morbidity and mortality in cryptococcal infections, particularly CNS infections. *C. neoformans* has a specific predilection for the CNS and can traverse the blood-brain barrier (BBB) through various mechanisms, including paracellular, transcellular, and Trojan horse pathways (11–14). Our previous RNA sequencing data on the host response to *C. neoformans* infection revealed the transcriptome of the human brain endothelium challenged with *C. neoformans* (15). In this study, we also analyzed the transcriptome of cryptococcal genes in response to human brain endothelial cells during transmigration across the blood-brain barrier (BBB) by comparing cryptococcal genes with and without incubation with human cerebral microvascular endothelial cells (hCMEC/D3) (Accession no. SRR32411784, SRR32411785). Notably, the cryptococcal *Amt* family of ammonium transporters was among the top three most highly upregulated genes in an in vitro BBB model, highlighting its potential role in this process. Previous studies from elsewhere indicated that the *Amt* gene family promotes an invasive phenotype characterized by pseudohyphal growth and facilitates dikaryon mating under ammonium-limiting conditions (16). Interestingly, *C. neoformans* can undergo morphological changes during direct transcytosis (17). Therefore, this study aims to investigate the role of the *Amt2* gene, a member of the *Amt* family of ammonium transporters, in the transmigration of *C. neoformans* across the blood-brain barrier (BBB) using an *in vitro* BBB model with hCMEC/D3. Additionally, the study seeks to determine whether the *Amt2* gene is associated with other common virulence factors produced by *C. neoformans*.

## Results

### The transmigration of *C. neoformans* across through the in vitro BBB model

In our previous study, the host’s transcriptome response to *C. neoformans* infection revealed that several cryptococcal genes were upregulated during *C. neoformans* transmigration, with the ammonium transporter gene (*AMT2*) showing a 1.98-fold increase (Figure 1).

**Figure 1.**
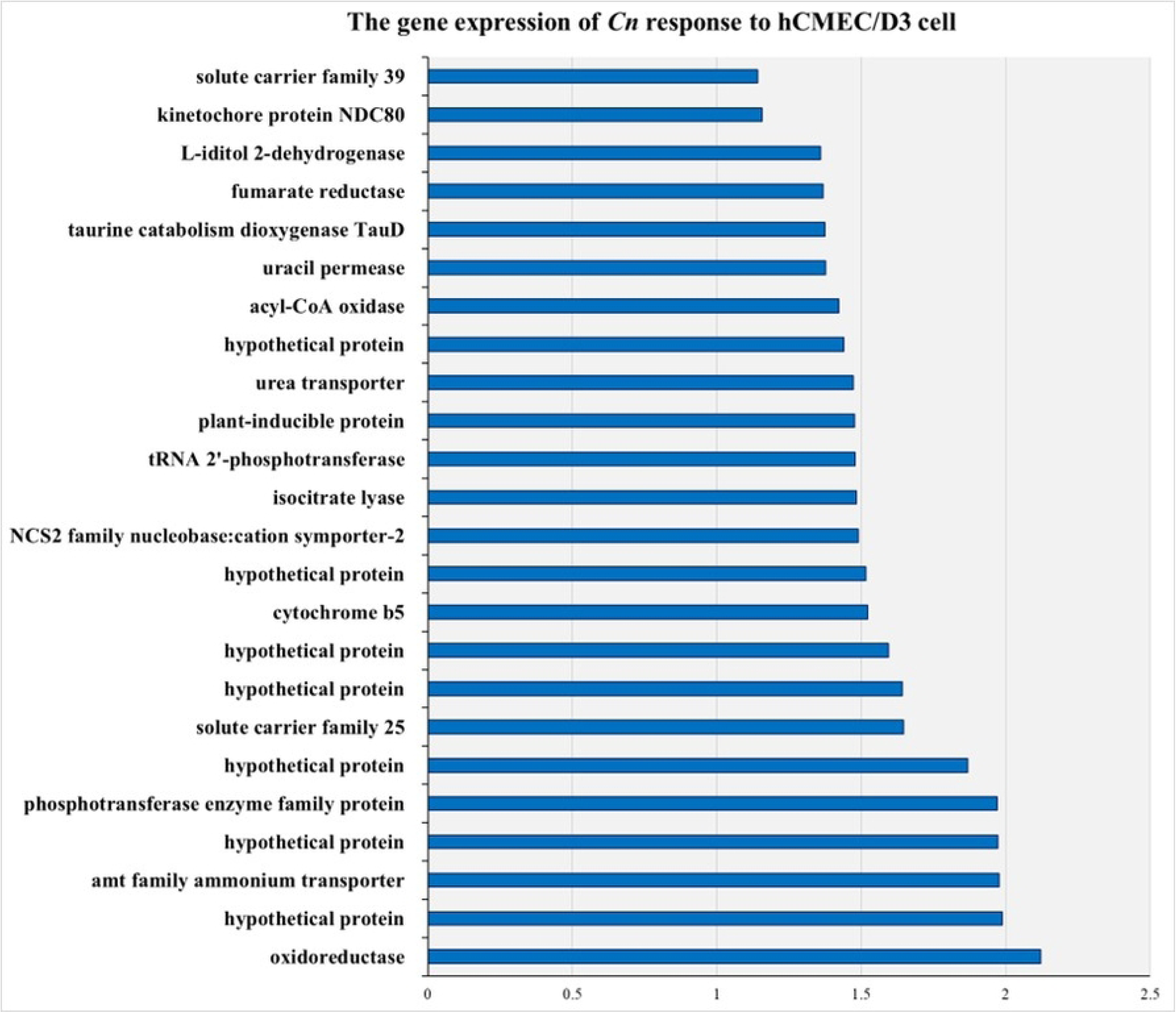
The gene expression profile of *C. neoformans* (H99) in response to human brain endothelial cells (hCMEC/D3).

Therefore, we investigated the association of this gene with the transmigration process and its relevance to other important virulence factors. We first compared the ability of the *amt2*Δ mutant strain to traverse the in vitro BBB transwell model (Figure 2A) using the hCMEC/D3 cell line. Before the transmigration assay, the TEER value was 48.03 ± 4.89 Ω·cm², confirming that the cells were fully confluent and had formed an intact barrier. The TEER value of the model of the endothelial monolayer cultured under static condition were around 30-50 Ω.cm² (18). The integrity of the BBB was confirmed by assessing monolayer membrane permeability using FITC-dextrans. The average FITC crossing ratio was 0.026 ± 0.01 before transcytosis and 0.082 ± 0.02 after transcytosis. In contrast, the negative control, which used a collagen-coated membrane without endothelial cells, exhibited a ratio of 0.97 ± 0.09 before transcytosis and 0.87 ± 0.02 after inoculation (Figure 2C). These results suggest that the tight junctions of the BBB model remained intact both before and after treatment with *C. neoform ans* in the transmigration assay.

**Figure 2.**
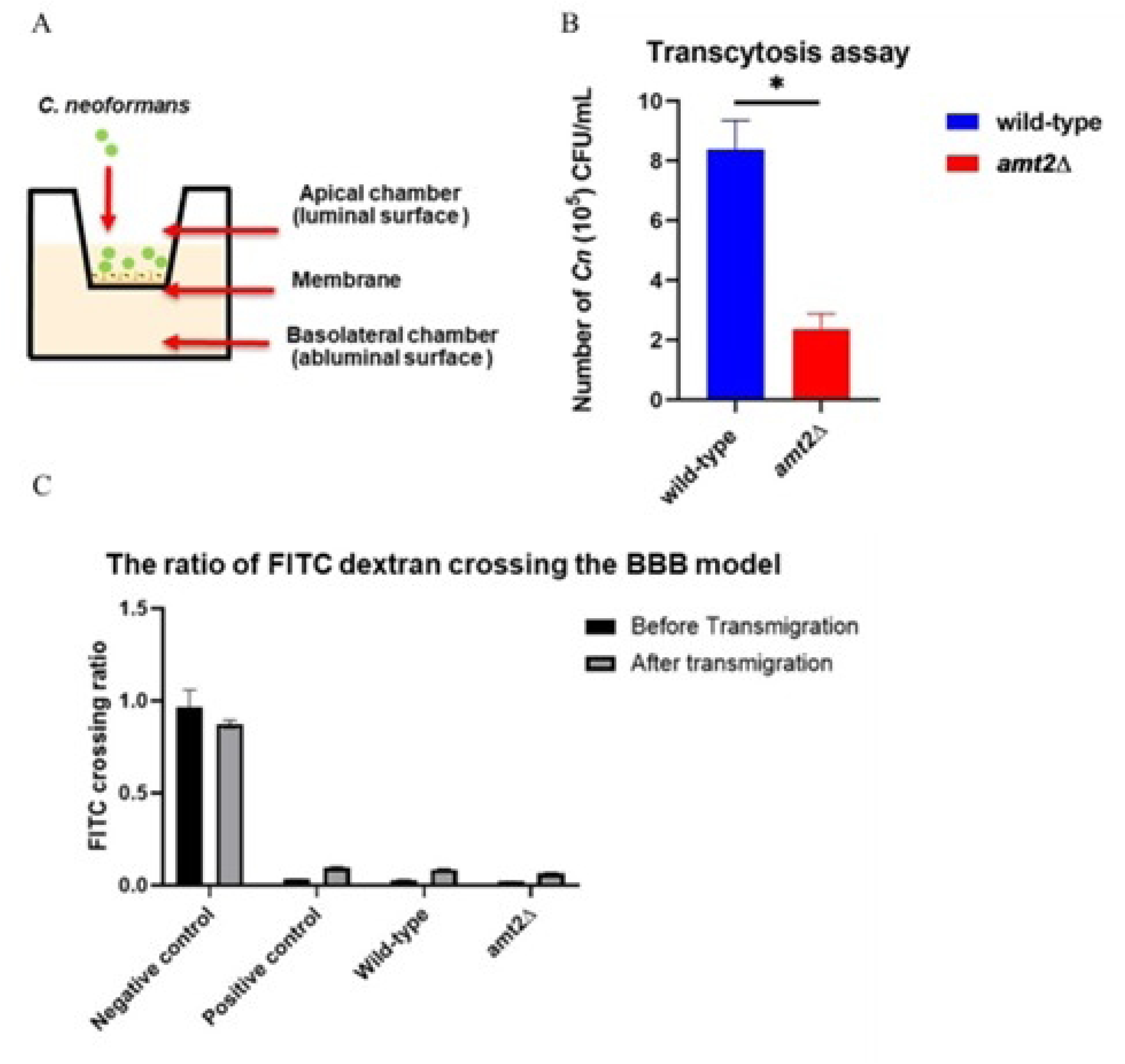
The *in vitro* model of BBB (A), The *in vitro* BBB model assay demonstrates that the *amt2*Δ mutant strain shows a defect in transmigration across the BBB when compared to the wild-type strain (B). The integrity of monolayer membrane, as assessed using FITC-dextrans, does not show a significant difference before and after treatment with *C. neoformans* (C). As analyzed by an unpaired t-test (Mean ± SEM, n = 3, \**p* < 0.05)

To assess whether the *amt2*Δ mutant strain can transmigrate through the endothelial cells, the suspension (yeast cells with EBM-2 media) collected from the abluminal side (lower chamber) was plated to determine colony-forming units (CFUs). The wild-type strain yielded 8.36 × 10^5^ ± 1.71 CFU/mL, whereas the *amt2*Δ mutant strain displayed a significantly reduced ability to transmigrate, with only 2.26 × 10^5^ ± 0.94 CFU/mL (Figure 2B). This reduction suggests that the *amt2*Δ strain exhibits impaired transmigration across the BBB.

### In vitro growth kinetics of yeast cells

To assess the growth kinetics, the knockout strain was cultured at two different temperatures: 30°C (environmental) and 37°C (mammalian host). The results showed that the exponential growth phase of the *amt2*Δ mutant strain was similar to that of the wild-type strain at both temperatures, with growth remaining stable over 2 to 3 days. No significant difference between the two strains was observed at the mammalian temperature, as determined by an independent t-test with *p* < 0.05 (Figure 3). These findings suggest that the *amt2*Δ mutant strain does not have a growth kinetics defect and is capable of growing and proliferating at 37°C, similar to the thermotolerant wild-type strain.

**Figure 3.**
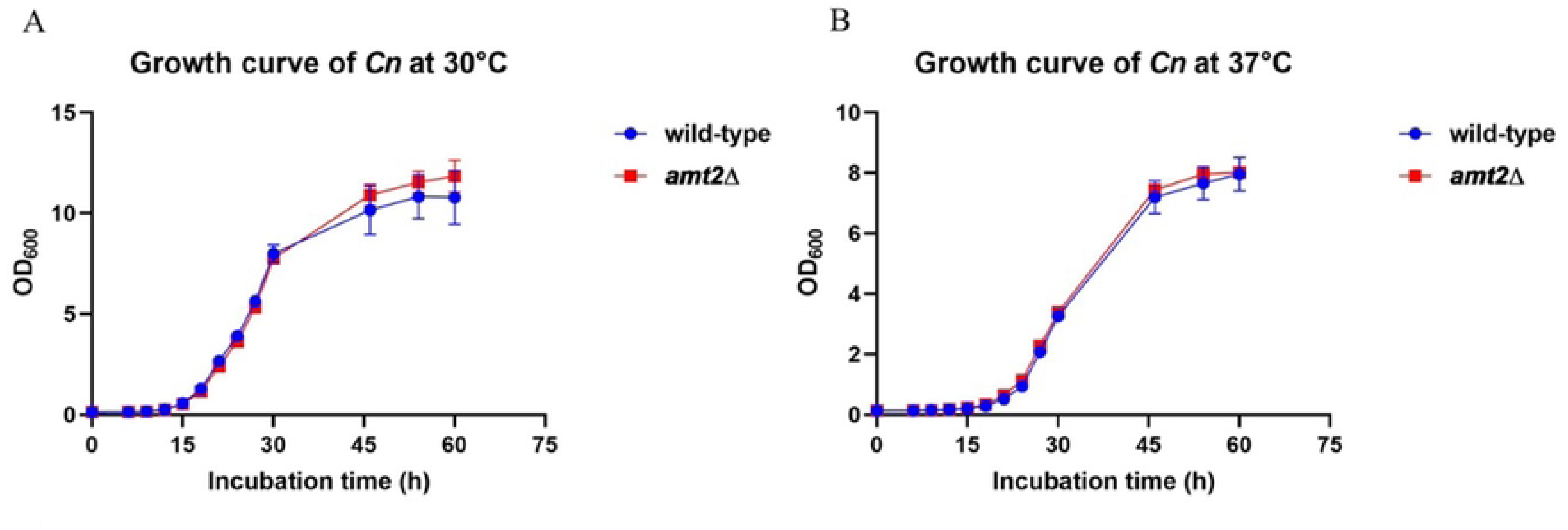

### Virulence factor of Cryptococcus neoformans

#### Capsule formation

Capsule formation was observed after 48 hours of incubation in DMEM with 20% FBS. The results showed that the polysaccharide capsule was slightly larger at 37°C compared to 30°C. Furthermore, the capsule size of the mutant strain was significantly smaller than that of the wild-type strain at both temperatures, as shown in Figure 4A. The average capsule sizes of the wild-type and *amt2*Δ mutant strains at 30°C were 1.63 ± 0.12 µm and 1.45 ± 0.14 µm, respectively. At 37°C, the wild-type strain exhibited a capsule size of 1.845 ± 0.15 µm, while the *amt2*Δ mutant showed a size of 1.61 ± 0.13 µm (Figure 4B-E). These findings suggest that deletion of the *Amt2* gene affects the synthesis of the polysaccharide capsule. Based on our results, the reduction in capsule size may be attributed to the loss of function of the *AMT2* transporter, which is involved in nitrogen transport into cryptococcal cells. Without the ability to efficiently transport nitrogen, a crucial factor for capsule formation, the signal for capsule production may be diminished.

**Figure 4.**
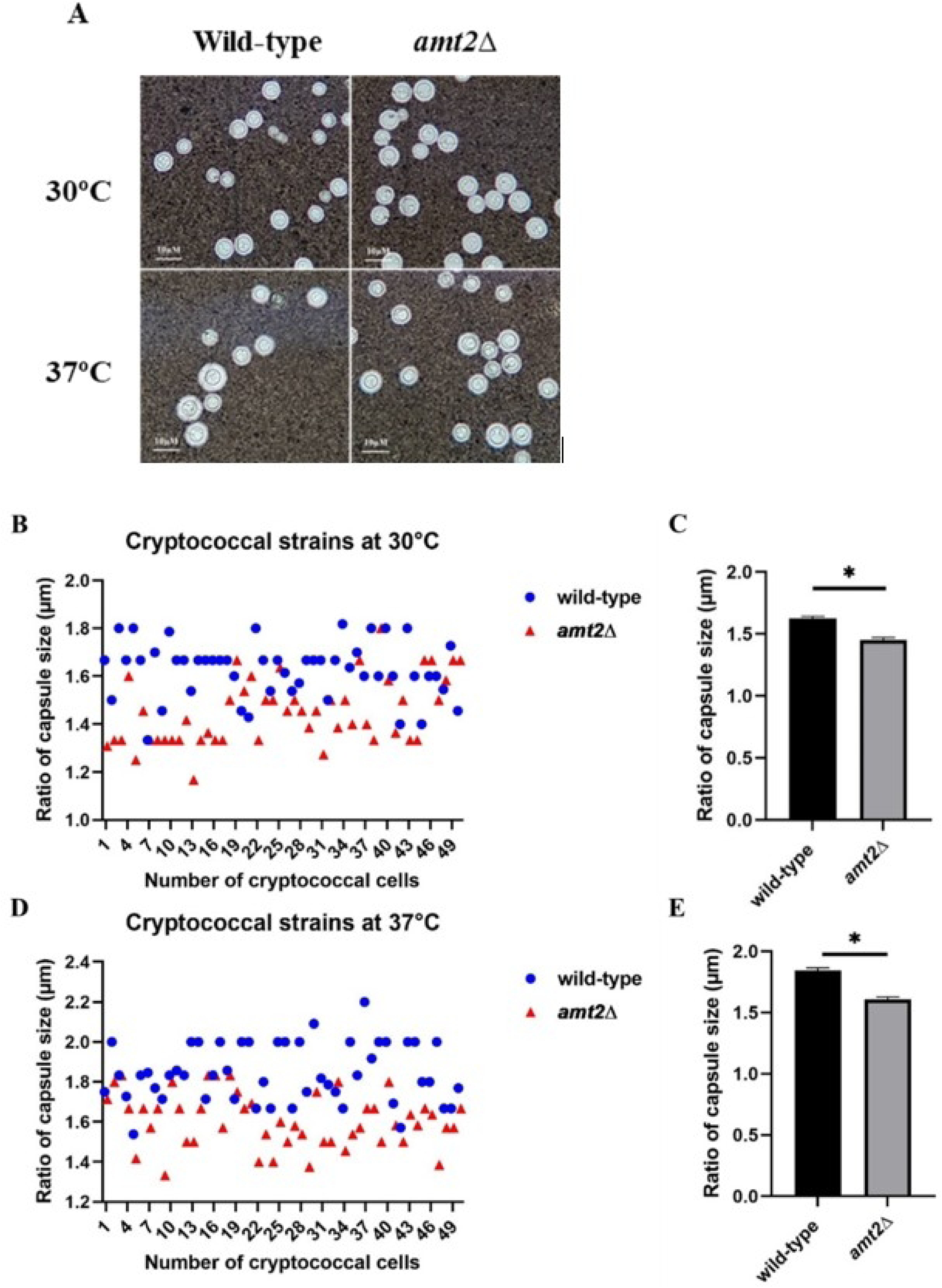
A comparison of the capsule sizes between both cryptococcal strains. The cryptococcal cells were stained with India ink and observed under a microscope (A). The capsule size ratios of the cryptococcal strains differed significantly at both 30°C (B-C) and 37°C (D-E). The *amt2*Δ mutant strain exhibited a smaller polysaccharide capsule compared to the wild-type strain, as determined by the Mann-Whitney U test (Mean ± SEM, n = 50, *p < 0.05).

### Urease activity

Our study evaluated urease production in the *amt2*Δ mutant strain using urea agar and RUH medium. On urea agar plates, the *amt2*Δ mutant strain demonstrated urease activity by hydrolyzing urea into ammonia, which caused the medium containing phenol red to change from light orange to pink. The results showed that the *amt2*Δ mutant strain could secrete urease at 3, 6, and 9 hours of incubation, although the reaction rate was slightly slower compared to the wild-type strain. The effects of different temperatures on the growth rate of the cryptococcal strains are shown in the colonies depicted in Figure 5A-B.

**Figure 5.**
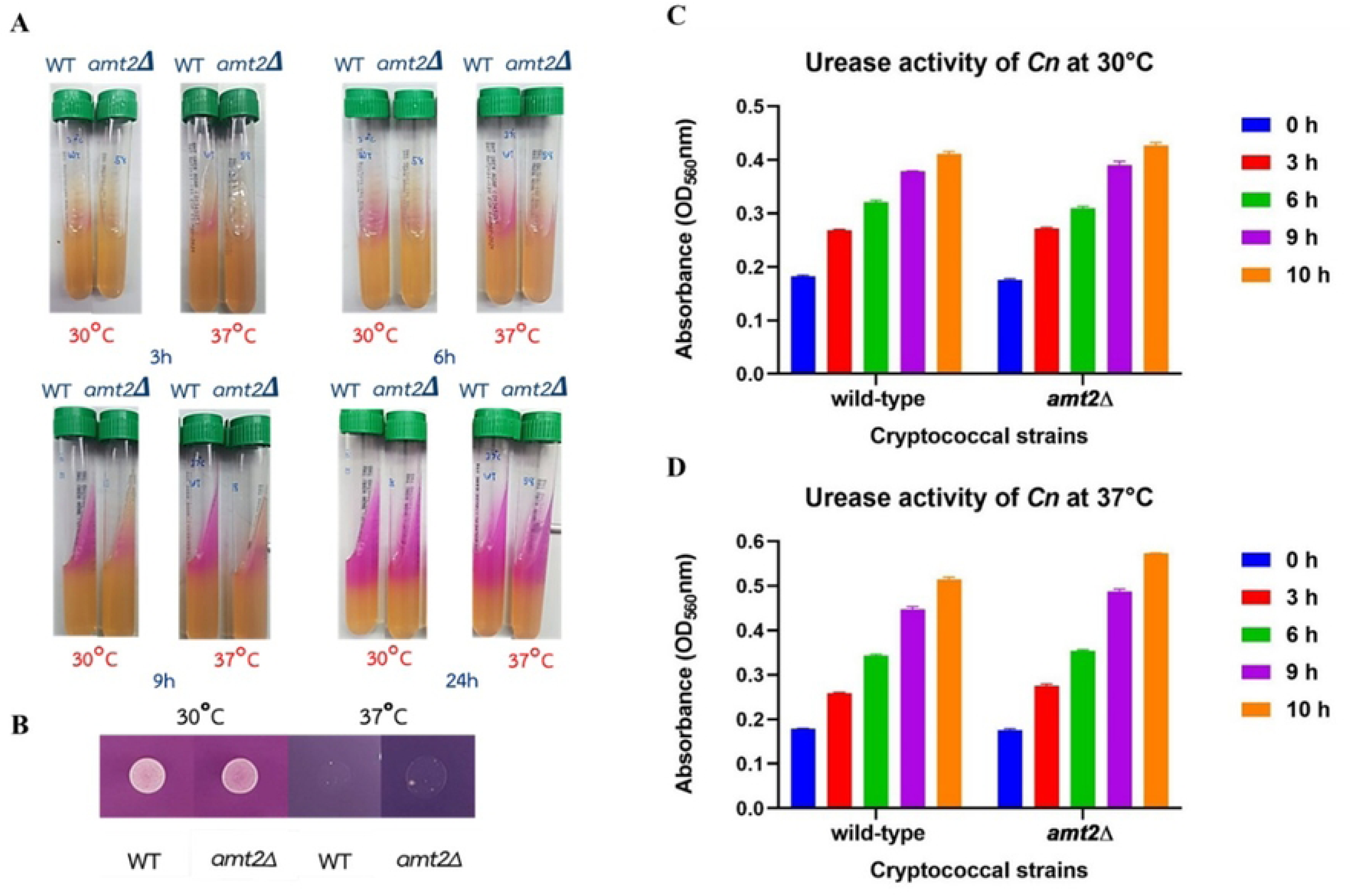
A comparison of the urease activity between both cryptococcal strains. The urease activity of the wild-type and *amt2*Δ mutant strains was evaluated on urea agar at 30°C and 37°C over 3, 6, 9, and 24 hours (A). At 48 hours, colony characteristics showed white colonies at 30°C, with better growth compared to incubation at 37°C (B). Ureolytic activity was further assessed using RUH broth at 3, 6, 9, and 10 hours, with the data analyzed using the Mann-Whitney U test (n = 3, p < 0.05) (C-D).

For quantitative analysis, ureolytic activity was assessed using RUH broth (results shown in Figure 5C-D. Both strains exhibited increased urease activity with longer incubation periods. No significant difference in urease levels was observed between the mutant and wild-type strains at the same incubation times. Thus, temperature variations did not affect the urease activity of either strain. Based on these findings, the absence of the *Amt2* gene did not significantly affect urease activity, suggesting that the mutant strain does not have a direct defect in urease production. Therefore, the *Amt2* gene may not be specifically involved in urease production in *C. neoformans*.

### Melanin production

CAFC test agar was used to identify and differentiate *C. neoformans* from other closely related yeast species based on its ability to produce melanin or melanin-like pigments. The results showed that both the wild-type and *amt2*Δ mutant strains formed brown or black colonies starting from day 2 of incubation at 37°C. Both strains demonstrated melanization by producing melanin or melanin-like pigments from the o-diphenol (caffeic acid) in the presence of ferric citrate. However, the mutant strain tended to produce more pigment, as evidenced by the darker black colonies compared to the wild-type strain after 4 days of incubation (Figure 6). This suggests that the *amt2*Δ mutant strain, lacking the *AMT2* gene, may experience stress, leading to increased melanin production, which serves as an antioxidant

**Figure 6.**
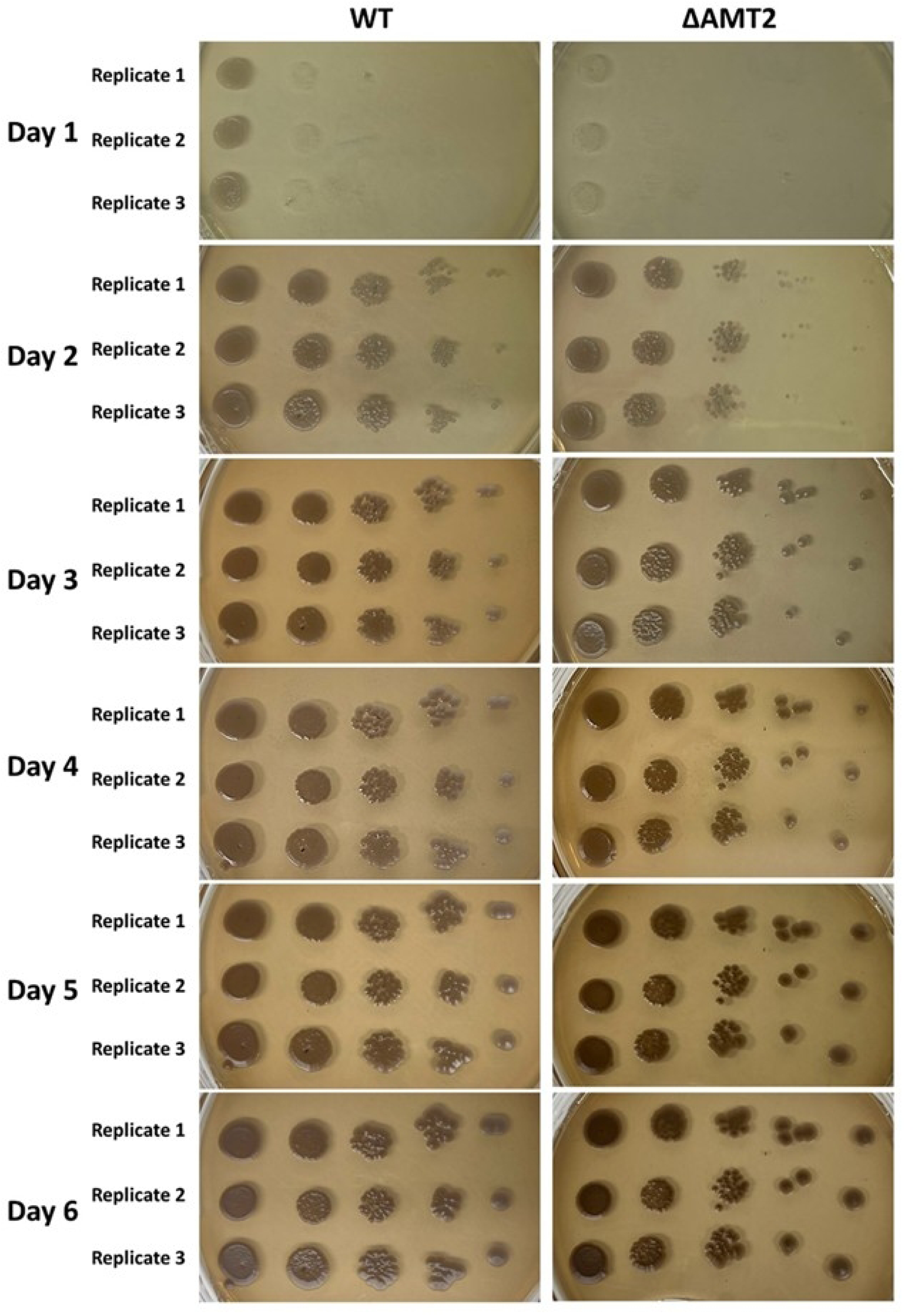
A comparison of the melanin production between both cryptococcal strains. Melanin production by the wild-type and mutant strains of *C. neoformans* was assessed on CAFC agar. The first spot represents the final inoculum adjusted to the 0.5 McFarland standard, with subsequent spots being five-fold dilutions. The plates were incubated at 37°C for 6 days.

### Phenotypic assay

Depending on environmental conditions, *C. neoformans* can undergo various developmental forms, such as yeast and filamentous types. Evidence suggests that *C. neoformans* can change its morphology during direct transcytosis. Consequently, we investigated whether the ammonium permease gene *Amt2* influences pseudohyphal development under nitrogen-limiting conditions and whether *Amt2* expression is sensitive to different nitrogen supply concentrations.

In this study, we observed that neither the wild-type nor the *amt2*Δ mutant strain formed pseudohyphae after 3, 5, and 7 days of incubation on YNB media containing 10 µM and 50 µM ammonium sulfate at both 30°C and 37°C, as shown in Figure 7A. Both the wild-type and mutant strains exhibited small, white colonies with a circular shape, flat surface, raised margins, and a mucoid texture. These results suggest that the *Amt2* gene did not trigger pseudohyphal growth, likely due to insufficient ammonium concentrations entering the cell to activate the ammonium transporter. Consistent with these findings, a phenotypic assay was performed using cryptococcal cells exposed to hCMEC/D3 cells. The results showed that neither the wild-type nor the *amt2*Δ mutant strain formed pseudohyphae after 3 hours. However, the cryptococcal cells retained their ability to internalize and continued to replicate through budding (Figure 7B). In conclusion, the deletion of the *Amt2* gene did not induce pseudohyphal growth in this study. However, the sensitivities of the mutant strain were similar to those of the wild-type strain, which may be attributed to the lack of a carbon source. Based on these findings, the in vitro phenotypic assay may not be suitable for comparing morphological changes between the wild-type and mutant strains.

**Figure 7.**
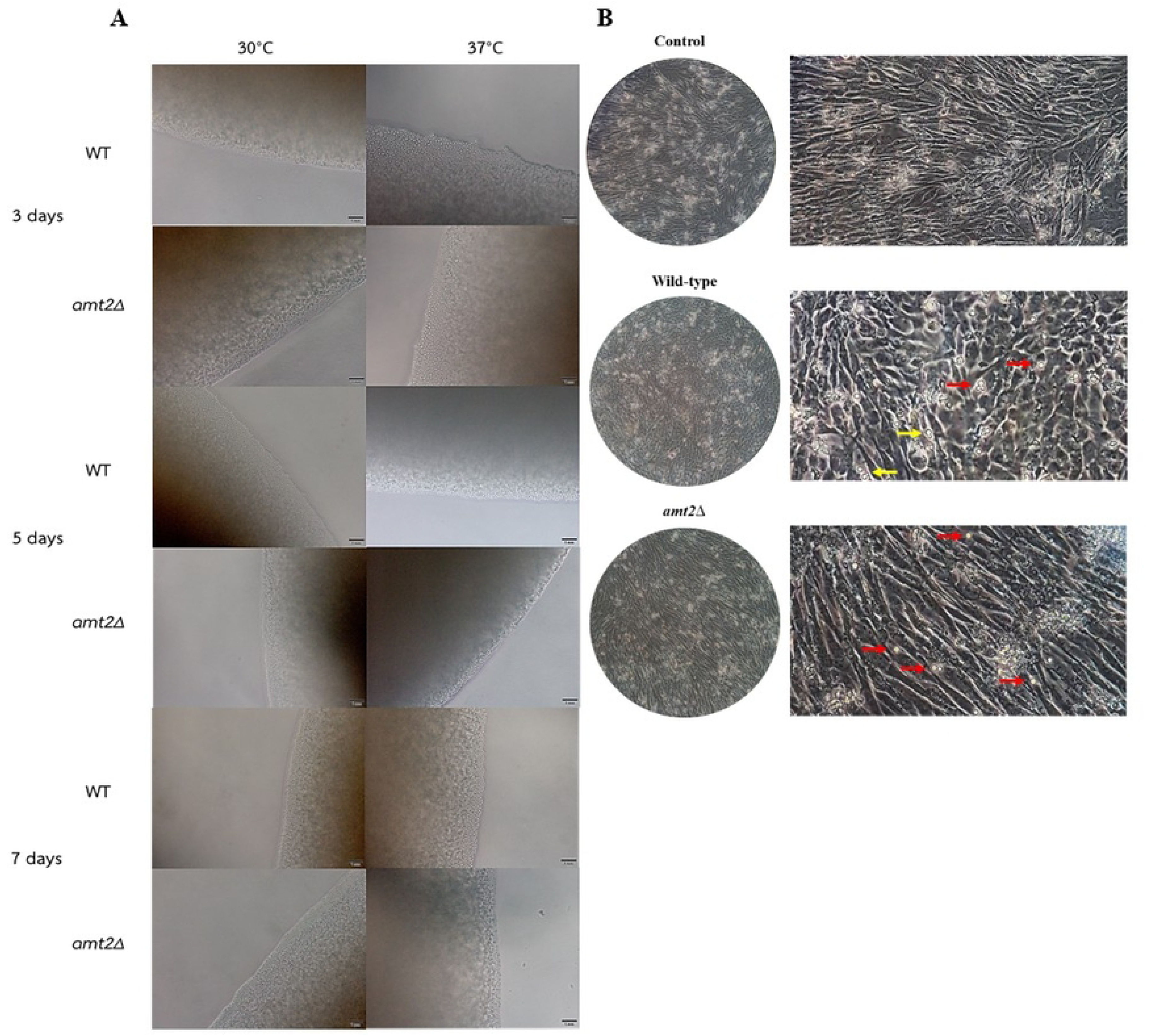
Morphology of cryptococcal cells in medium culture and cell culture. Phenotypic assay results after incubation for 3, 5, and 7 days on YNB media were observed under a light microscope with a 20X objective lens (A). In the phenotypic assay of cryptococcal cells exposed to hCMEC/D3 cells after 3 hours of incubation, neither strain formed pseudohyphae. However, cryptococcal cells were internalized by brain endothelial cells (red arrow), and *C. neoformans* propagated through budding (yellow arrow) (B).

### The sensitivity of *C. neoformans* growth under the limiting-nitrogen source

This study utilized a sensitivity assay to assess growth rates in response to different nitrogen sources, specifically evaluating *Amt2* activity. The growth rate did not differ significantly between the wild-type and mutant strains on media containing ammonium concentrations of 10 µM, 30 µM, and 50 µM (Figure 8). Both strains also exhibited similar colony morphology, characterized by a circular shape, convex elevation, and small white colonies. These results suggest that the absence of the *Amt2* gene did not affect *C. neoformans* ability to utilize nitrogen sources, likely due to the compensatory activity of the *Amt1* gene.

**Figure 8.**
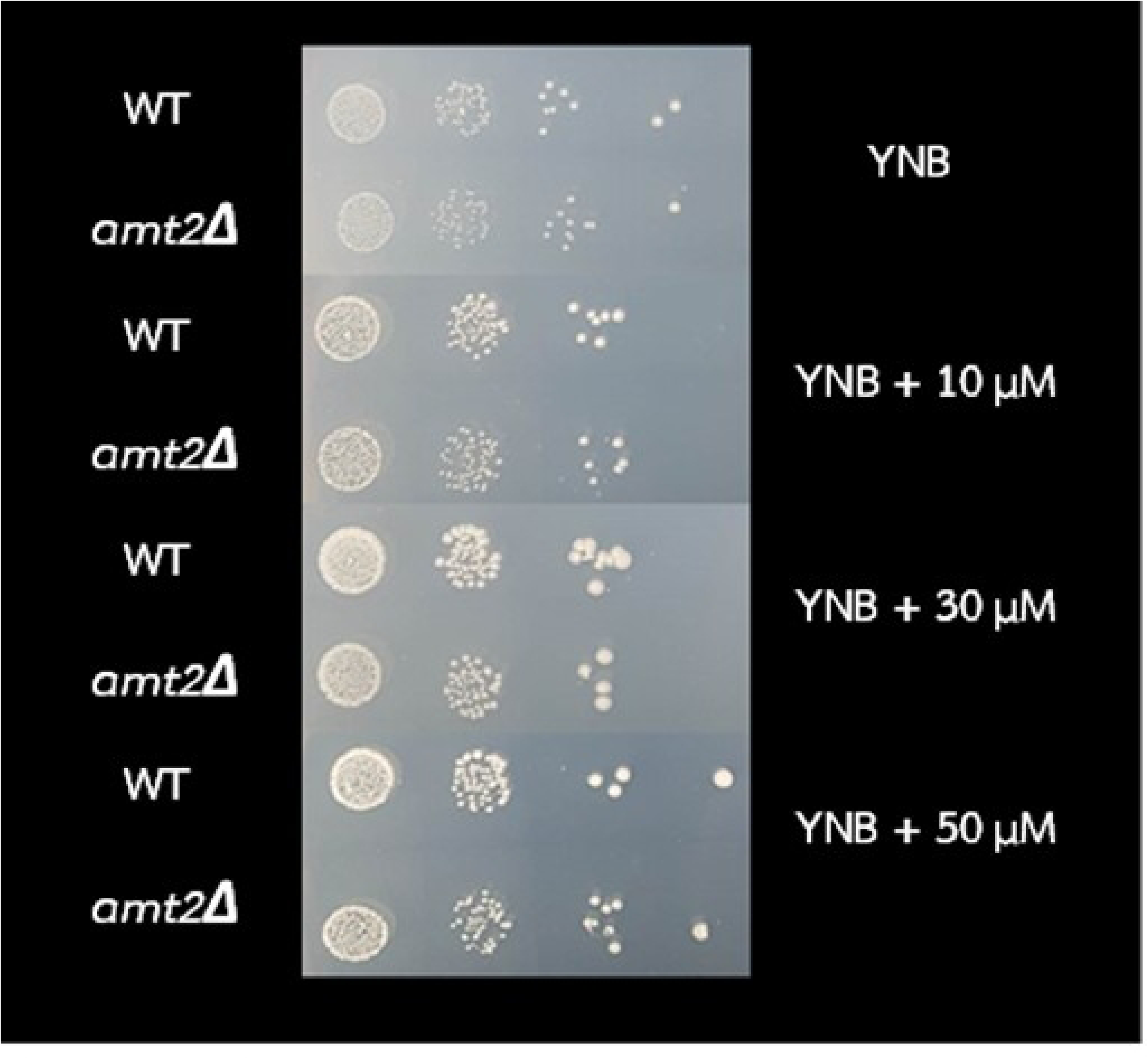
Sensitivity of cryptococcal strains to limiting concentrations of ammonium. The sensitivity assay for *C. neoformans* was performed by serially diluting the wild-type and *amt2*Δ mutant strains, then spotting them onto media with varying ammonium concentrations (10 µM, 30 µM, and 50 µM).

## Discussion

*C. neoformans* is a pathogenic encapsulated yeast that causes central nervous system infections through various mechanisms, including paracellular, transcellular, and Trojan horse pathways. In this study, we investigated the pathway *C. neoformans* uses to traverse brain endothelial cells and found that it primarily employs the transcellular route. This process likely involves the interaction of endothelial cell molecules with the pathogen, facilitating enhanced transmigration while maintaining the integrity of the blood-brain barrier (BBB).

Several key factors have been identified that contribute to the mechanisms underlying the transcellular process by which *C. neoformans* crosses the blood-brain barrier (BBB). Reports suggest that the glycoprotein CD44, expressed by brain microvascular endothelial cells (BMECs), enhances the binding of *C. neoformans* to hyaluronic acid. This interaction triggers rearrangement of the filamin and actin cytoskeleton, which is linked to the *CPS1* gene (19). Additionally, EphA2 activity via CD44 creates a permeable barrier that facilitates the migration of *C. neoformans* across the blood-brain barrier (BBB) (15). The novel metalloprotease MPR1 is also required for the attachment and internalization of *C. neoformans* by brain endothelial cells without disrupting protein complexes (8). Moreover, genes such as *PLB1*, *ITR1*a, *ITR3*c, and *RUUB1* have been implicated in the transcellular traversal of *C. neoformans* through the brain (20–22). Real-time imaging of transmigration across the BBB revealed that *C. neoformans* becomes arrested in brain capillaries and undergoes a morphological change to an oval shape during brain infection. This transformation is consistent with how the fungus adapts to the brain environment (17).

Our previous RNA sequencing data, which examined the overall transcriptome of the wild-type *C. neoformans* (H99) during transmigration through microvascular endothelial cells (15), provided insights into the transcriptome of the human brain endothelium in response to *C. neoformans*. In the current study, we present the transcriptomic profile of *C. neoformans* in response to the human brain endothelium. Notably, we identified ammonium transporter genes as among the top three most upregulated genes in *C. neoformans* during this process. These proteins belong to the Amt/Mep/Rh family of permeases commonly found in pathogenic fungi. The *Amt* (ammonium permeases) and *Mep* (methylammonium permeases) genes were first cloned from *Arabidopsis thaliana* (23) and *Saccharomyces cerevisiae* (24), respectively. In *S. cerevisiae*, MEP proteins play a crucial role in both nitrogen transport and sensing. Specifically, *Mep2* is involved in sensing low ammonium availability and is linked to signal transduction pathways that lead to pseudohyphal or filamentous growth (25, 26).

In our recent study, we found that the *C. neoformans* H99 strain lacking the *Amt2* gene exhibits a defect in transmigration across the blood-brain barrier (BBB), emphasizing the crucial role of the *Amt2* gene in BBB traversal. Our findings suggest that *C. neoformans* penetrates the BBB via a transcellular mechanism, as the integrity of the brain endothelial cells remained largely intact after infection with the cryptococcal strains. Deletion of the *Amt2* gene in *C. neoformans* reduced its ability to transmigrate, thereby impairing its capacity to cause cryptococcal infection. Additionally, the *amt2*Δ mutant displayed reduced capsule formation compared to the wild-type H99 strain. However, no morphological differences were observed between the wild-type and mutant strains in either the culture medium or the BBB model. Regarding capsule formation, it has been noted that capsule size increases during the stationary phase (around 48 hours), compared to the exponential growth phase (27). Our results suggest that the deletion of the *Amt2* gene affects capsule size in the mutant strain. Nitrogen sources play a critical role in inducing capsule production in *C. neoformans*, with nitrogen metabolism serving as a key signal for capsule induction (28). If *C. neoformans* loses its ability to transport nitrogen, it could result in a diminished signal for capsule production in the mutant strain. A reduction in capsule size is linked to increased phagocytosis by the host immune response. Studies have shown that a larger capsule can protect yeast cells from host immune defenses by inhibiting phagocytosis, leukocyte migration, and cytokine production (29). Mandal, R., et al reported in vivo infections resulting in the different sizes of capsules formation, with the lung and brain displaying a more pronounced response compared to other organs (30). Capsule biosynthesis is directly regulated by capsule-associated genes (*CAP* genes), such as *CAP10*, *CAP59*, *CAP60*, and *CAP64* (26). Although many genes are involved in capsule biosynthesis, the regulation of nitrogen metabolism also affects the capsule size in *C. neoformans*.

The growth ability of both the wild-type and mutant strains at 37°C was comparable, indicating that the observed defect in the transcytosis assay is not due to any growth disadvantage of the mutant strain at this temperature. Additionally, the mutant strain exhibited slightly slower urease production compared to the wild-type strain. This reduced urease activity may contribute to the diminished efficiency of the mutant strain during the early stages of the transmigration process, although the mutant still retains ureolytic activity. Urease plays a crucial role in *C. neoformans* ability to cross the blood-brain barrier (BBB) by converting urea into ammonia. This process may facilitate migration through paracellular pathways or trigger signal transduction pathways that induce morphological changes (8, 14). Urea is a byproduct of protein metabolism and presents in large amounts in cerebrospinal fluid (CSF) (31). The urease enzyme catalyzes the hydrolysis of urea, resulting in the production of ammonia (NH₄⁺) and carbonic acid (H₂CO₃) a critical virulence factor that neutralizes the acidic microenvironment (32). This process helps pathogens survive the harsh pH conditions within phagolysosome (33). *C. neoformans* has the ability to form pseudohyphae during host infection through the activity of *Amt* genes, including *Amt1* and *Amt2*, particularly under ammonium-limiting conditions (16). However, our results showed that neither the wild-type nor the *amt2*Δ mutant strain exhibited morphological changes. Previous studies have demonstrated pseudohyphal growth in *C. neoformans* when cultured on YNB medium supplemented with 2% glucose as the nitrogen source (34). The carbon source plays a pivotal role in fungal development, often driving a morphological transition from budding to pseudohyphal growth (35). During infections, the primary carbon sources available in the host include glucose, lactate, and acetate (36). Moreover, unisexual mating in *C. neoformans* has been shown to trigger pseudohyphal growth at the colony edges when grown on media with fermentable carbon sources, such as glucose, galactose, or sucrose. As a result, the experimental conditions in our study may not have been optimal for inducing pseudohyphal formation in the cryptococcal strains. The morphological changing in C. *neoformans* with enlarged titan cells and smaller cells alters the immune response and facilitates the spread from the lungs to other organs (37).

Our current study demonstrated that the deletion of *Amt2* alone did not affect the sensitivity of *C. neoformans* to nitrogen availability or its ability to undergo pseudohyphal growth under low-ammonium conditions. However, previous evidence shows that the double mutant strain, *amt1*Δ*amt2*Δ, exhibited increased sensitivity and was unable to form pseudohyphae under similar low-ammonium conditions (38). In conclusion, the growth abilities of the wild-type and *amt2*Δ mutant strains were comparable, suggesting that *C. neoformans* can produce or secrete the metabolites necessary for growth. These metabolites are likely associated with its virulence factors, which may contribute to the fungus’s ability to traverse the blood-brain barrier, playing a key role in cryptococcal infection. Evidence suggests that the double mutant strain lacking both *Amt1* and *Amt2* failed to produce pseudohyphae and instead formed smooth colonies. While *Amt1* and *Amt2* share similar functions, *Amt2*-deficient strains did not exhibit invasive growth and showed reduced mating under low ammonium conditions. In contrast, *Amt1*-deficient strains maintained levels of invasive growth comparable to the wild-type (38). These findings imply that pseudohyphal formation requires limited, rather than absent, carbon and nitrogen sources. A deficiency in both nutrients seems to promote the development of diploid yeast and pseudohyphal growth. In both environmental and host settings, *C. neoformans* can produce melanin pigments from various phenolic compounds, including catecholamines and nitrogen-containing diphenolic compounds, which serve as precursors (39). Laccase activity was observed in both the wild-type and mutant strains, with the mutant strain showing stronger melanin production. This increased production may result from the absence of *AMT2*, which induces stress in the mutant strain and triggers enhanced melanin synthesis as a protective mechanism against oxidative stress. *C. neoformans* synthesizes brown-colored eumelanin through the action of laccases (*Lac1* and *Lac2*) which is low expression under nutrient-rich conditions, but expression is upregulated under nutrient starvation (40, 41).

In conclusion, the *Amt2* gene plays a crucial role in the transmigration of *C. neoformans* through the blood-brain barrier (BBB). The observed defect in the mutant strain lacking *Amt2* may be linked to its slower urease activity, although the precise mechanisms underlying this defect warrant further investigation. Additionally, the *Amt2* gene is essential for capsule formation in *C. neoformans*, though it does not appear to impact the yeast’s ability to grow at host temperature. Melanin production is enhanced in the mutant strain, likely offering increased protection against oxidative stress. Furthermore, the *Amt2* gene alone does not seem to influence pseudohyphal development, possibly due to the absence of suitable carbon sources in the tested conditions. Our study has identified a critical gene involved in the transmigration process of *C. neoformans*, which could provide valuable insights for developing future therapeutic strategies, including drug development, to prevent cryptococcal infections. Further research into the pathways involving *Amt2* and its interactions with other molecules is essential to better understand how this gene contributes to the transmigration and infection processes of *C. neoformans*.

## Materials and Methods

### Fungal strains

*C. neoformans* strain H99 (serotype A) and *amt2Δ* mutant strain (04758; Ammonium permease gene) were obtained from the Hiten Madhani lab (UCSF, USA). The fungal strains were freshly retrieved from -80°C frozen stock prior to use in the experiment. They were plated on Yeast Extract-Peptone-Dextrose (YPD) agar and incubated at 30°C for 72 hours. The resulting *C. neoformans* colonies were then grown in YPD broth at 30°C with agitation for 18 hours.

### hCMEC/D3 cells and culture conditions

The human cerebral microvascular endothelial cell line, hCMEC/D3, was purchased from the American Type Culture Collection (ATCC). The cells were seeded in a collagen-type I-coated culture flask and grown to confluence in EBM-2 medium with growth factors (EGM-2 MV Bullet Kit, Lonza, Walkersville, MD, USA) at 37°C in a 5% CO_2_ atmosphere.

### Transcytosis assay in the in vitro model of BBB

The hCMEC/D3 cells, with fewer than 35 passages, were used for the transcytosis assay. The in vitro static monolayer model featured a trans-well apparatus with an upper (luminal) chamber and a lower (abluminal) chamber, divided by a collagen-coated porous membrane (8 µm; Corning). 1 x 10^4^ hCMEC/D3 cells were seeded on the upper side. During incubation, the medium was replaced every 3-4 days, gradually decreasing from 1x strength to 0.5x, and then to 0.25x. The integrity of the monolayer was assessed using the trans-endothelial electrical resistance (TEER) measurement assay. Additionally, the integrity of the BBB model was confirmed before the transcytosis assay using a permeability assay. Cryptococcal cells (1 x 10^6^) were added to the upper chamber and incubated overnight, after which the migrated cells from the lower chamber were collected for colony-forming unit (CFU) determination.

### FITC/Dextran permeability assay

FITC-conjugated dextran (relative molecular mass, 70K) assessed permeability across the endothelial cell monolayer. FITC/Dextran was used in parallel for the transcytosis assay. Before and after the transcytosis assay of *C. neoformans*, hCMEC/D3 cells permeability was measured by adding 1 mg/ml of FITC-labeled dextran and measured for fluorescence at 538 nm when excited at 485 nm with a spectrophotometer (Varioskan Lux multimode microplate reader scientific, Singapore). Each upper and bottom chamber sample was read in triplicates on a fluorescent plate (Thermo Scientific, USA). The FITC/dextran ratio was calculated by measuring the fluorescence intensity in the bottom chamber relative to that in the upper chamber. The statistical analysis was performed using Prism software. Statistical analysis, at *P*-value <0.05, was considered significantly different.

### *In vitro* growth kinetics of yeast and fungal cells

Cryptococcal strains were cultured in YPD broth and incubated with shaking at 200 rpm for 24 hours at 30°C and 37°C. The starting inoculum’s optical density was set to 0.1 at 600 nm (OD_600_). Using a spectrophotometer, OD_600_ measurements were taken every 3 hours for the first 24 hours and then every 6 hours until 72 hours. The experiments were conducted in triplicate.

### Capsule formation

A single colony from YPD agar of each strain was inoculated into YPD broth and incubated overnight at 30°C. The resulting suspensions were washed and resuspended in 1X PBS to an OD600 of 0.7. These were then incubated in 2 mL of DMEM (Dulbecco’s Modified Eagle’s Medium with 20% Fetal Bovine Serum) at 30°C and 37°C for 48 hours with shaking. After incubation, the samples were centrifuged at 5,500 rpm for 3 minutes and resuspended in 500 µL of 1X PBS. Cell suspensions were mixed with Indian ink to visualize capsules and examined under a light microscope using a 40x objective lens. At least ten randomly chosen fields were photographed, and 50 cells were analyzed.

### Urease activity

After 18 hours of growth in YPD broth, cryptococcal cells were centrifuged and resuspended in 1X PBS. A 5 µL aliquot of the suspension (OD_600_ of 0.7) was streaked onto urea agar and spotted onto urea tubes. Both methods were incubated at 30°C and 37°C and observed periodically during the incubation. For additional urease activity analysis, RUH broth was used. Cryptococcal cells were incubated with rotation at 200 rpm at 30°C and 37°C for 10 hours. Suspensions were collected at 3, 6, 9, and 10 hours, centrifuged at 5,500 rpm, and measured at OD_560_ nm using a spectrophotometer. The assay was performed in triplicate for each time point.

### Melanin production

The wild-type and mutant strains of *C. neoformans* were tested for melanin production on Caffeic Acid Ferric Citrate (CAFC) agar (42). Each isolate was inoculated into YPD broth at 37°C for 24 hours. The cultured broths were cross-streaked on YPD agar and incubated at 37°C for 48 hours. Pure colonies were selected and prepared to a 0.5 McFarland standard in a 0.85% NaCl solution. The cell suspensions were then serially diluted in a five-fold series. Samples from each dilution (5 µl) were plated in triplicate on CAFC agar. The CAFC plates were incubated at 37°C for 6 days, and melanin production was indicated by the formation of black pigment colonies.

### In vitro phenotypic assay

The extent of invasive growth of cryptococcal strains was assessed under nitrogen-limiting conditions. The wild-type and mutant strains were grown on Yeast Nitrogen Base (YNB) media, which lacked amino acids and contained 10 µM, 30 µM, and 50 µM ammonium sulfate ((NH_4_)2SO_4_) at 30°C and 37°C for one week. Invasive growth was evaluated by examining the agar surface under a microscope to identify pseudohyphae formation. This experiment was conducted in triplicate.

### The sensitivity assay

The sensitivity of *C. neoformans* to nitrogen sources was tested on a YNB medium containing varying ammonium concentrations (10 µM, 30 µM, and 50 µM). Cryptococcal strains were adjusted to an equal cell number, and 5 µL of each suspension was spotted onto YPD agar supplemented with different concentrations of (NH_4_)_2_SO_4_. Serially diluted cell suspensions were used for this assessment.

### Statistical analysis

Data are presented as mean ± standard error of the mean (SEM) from at least three independent experiments. Statistical significance was determined with \**p* < 0.05 for wild-type and *amt2*Δ mutant strains comparisons. An unpaired t-test was used to compare two groups in the transcytosis assay, while the Mann-Whitney U test was employed for virulence factor assays using GraphPad Prism (GraphPad Software, Boston, MA).

## Data Availability Statement

RNA Sequence data that support the findings of this study have been deposited in the National Center for Biotechnology Information (NCBI) with the primary accession code SRR32411784 and SRR32411785.

## Acknowledgements

We would like to acknowledge the Ministry of Higher Education, Science, Research and Innovation, Thailand, for the funding (Grant no. RGNS 64-120). We also thank the Center for Scientific and Technological Equipment, Suranaree University of Technology, the Thammasat University Research Unit in Nutraceuticals and Food Safety, and the Research Group in Medical Biomolecules, Faculty of Medicine, Thammasat University. I would also like to thank Aggie Gelli (Gelli Lab) for her valuable suggestions and insightful discussion on the project concept.

## Author contributions statement

**Conceptualization**: Sainamthip Rangdist, Ekkasit Kanklang, Methee Chayakulkeeree, Mantana Jamklang.

**Data Curation**: Sainamthip Rangdist, Ekkasit Kanklang, Natchaya Pakdeesiriwong

**Formal Analysis**: Sainamthip Rangdist, Ekkasit Kanklang, Natchaya Pakdeesiriwong, Watsana Penkhrue, Chompunoot Wangboon.

**Funding Acquisition:** Mantana Jamklang.

**Investigation:** Sainamthip Rangdist, Ekkasit Kanklang, Natchaya Pakdeesiriwong.

**Methodology**: Oratai Weeranantanapan, Sainamthip Rangdist, Ekkasit Kanklang, Natchaya Pakdeesiriwong.

**Project Administration**: Mantana Jamklang.

**Resources**: Oratai Weeranantanapan, Pathanin Chantree, Pongsakorn Martviset, Sirilak Chumkiew, Mantana Jamklang.

**Software**: Sainamthip Rangdist, Ekkasit Kanklang, Mantana Jamklang.

**Supervision**: Methee Chayakulkeeree, Mantana Jamklang.

**Validation**: Pongsakorn Martviset, Methee Chayakulkeeree, Mantana Jamklang.

**Visualization** Sainamthip Rangdist, Ekkasit Kanklang, Pongsakorn Martviset, Mantana Jamklang.

**Writing – Original Draft Preparation**: Sainamthip Rangdist, Ekkasit Kanklang, Pongsakorn Martviset, Mantana Jamklang.

**Writing – Review & Editing**: Sainamthip Rangdist, Ekkasit Kanklang, Natchaya Pakdeesiriwong, Watsana Penkhrue, Chompunoot Wangboon, Oratai Weeranantanapan, Sirilak Chumkiew, Pathanin Chantree, Pongsakorn Martviset, Methee Chayakulkeeree, and Mantana Jamklang.

